# Hybrid metachronal rowing augments swimming speed and acceleration via increased stroke amplitude

**DOI:** 10.1101/2021.04.22.441008

**Authors:** Mitchell P. Ford, William J. Ray, Erika M. DiLuca, S. N. Patek, Arvind Santhanakrishnan

**Affiliations:** School of Mechanical and Aerospace Engineering, Oklahoma State University, Stillwater, OK 74078, USA; Department of Biology, Duke University, Durham, NC 27708, USA

**Keywords:** Metachronal, Swimming, Rowing, Crustacean, Mantis shrimp

## Abstract

Numerous aquatic invertebrates use drag-based metachronal rowing for swimming, in which closely spaced appendages are oscillated starting from the posterior, with each appendage phase-shifted in time relative to its neighbor. Continuously swimming species such as Antarctic krill generally use “pure metachronal rowing” consisting of a metachronal power stroke and a metachronal recovery stroke, while burst swimming species such as many copepods and mantis shrimp typically use “hybrid metachronal rowing” consisting of a metachronal power stroke followed by a synchronous or nearly synchronous recovery stroke. Burst swimming organisms need to rapidly accelerate in order to capture prey and/or escape predation, and it is unknown whether hybrid metachronal rowing can augment acceleration and swimming speed compared to pure metachronal rowing. Simulations of rigid paddles undergoing simple harmonic motion showed that collisions between adjacent paddles restrict the maximum stroke amplitude for pure metachronal rowing. Hybrid metachronal rowing similar to that observed in mantis shrimp (*Neogonodactylus bredini*) permits oscillation at larger stroke amplitude while avoiding these collisions. We comparatively examined swimming speed, acceleration, and wake structure of pure and hybrid metachronal rowing strategies by using a self-propelling robot. Both swimming speed and peak acceleration of the robot increased with increasing stroke amplitude. Hybrid metachronal rowing permitted operation at larger stroke amplitude without collision of adjacent paddles on the robot, augmenting swimming speed and peak acceleration. Hybrid metachronal rowing generated a dispersed wake unlike narrower, downward-angled jets generated by pure metachronal rowing. Our findings suggest that burst swimming animals with small appendage spacing, such as copepods and mantis shrimp, can use hybrid metachronal rowing to generate large accelerations via increasing stroke amplitude without concern of appendage collision.

## Introduction

Coordinated rowing of multiple appendages is a common locomotion strategy in numerous aquatic organisms (Schabes and Hamner 1992; Walker 2002; Lim and DeMont 2009; Alben et al. 2010; Murphy et al. 2011; Colin et al. 2020). Metachronal rowing, where appendages are oscillated in sequence from the posterior to anterior with a time delay (i.e., non-zero phase lag) between neighbors, has been shown to significantly increase swimming speed for closely spaced appendages as compared to synchronous rowing with zero phase lag (Alben et al. 2010; Ford and Santhanakrishnan 2021a). Despite the broad morphological diversity across organisms that use metachronal rowing, the ratio of appendage spacing (*G*) to appendage length (*L*) across a number of taxonomically diverse species has been reported to fall in a narrow range from 0.2 to 0.7 (Murphy et al. 2011). Our recent study showed that smaller appendage spacing ratio (*G/L*) promotes hydrodynamic interactions that can increase swimming speed, and that stroke amplitude (*θ*) affects swimming speed more than phase lag for a given *G/L* (Ford and Santhanakrishnan 2021b). Decreasing *G/L* to augment swimming speed, by either increasing *L* or decreasing *G*, requires a reduction of *θ* to prevent appendage collision (or interference). To prevent appendage collisions, the maximum allowable *θ* is also limited by the phase lag, as increasing the phase lag would bring the tips of neighboring appendages in closer proximity at select instances in a metachronal stroke. It is thus necessary to balance the benefits gained from increasing *θ* with the benefits gained from increasing phase lag. Appendage flexibility can reduce the risk of damage from colliding paddles operating at or near the maximum *θ*, but flexible appendages are still limited to that stroke amplitude. For a fixed body length, *G/L* and number of appendages, stroke kinematics need to be altered to increase *θ* for augmenting swimming speed.

Animals have evolved multiple ways to modify stroke kinematic parameters and increase *θ*. One strategy is to vary the mean stroke angle of each appendage along the body length, such that appendages near the anterior end have a more anterior orientation and more posterior appendages have a more posterior orientation (Murphy et al. 2011). Mysids vary the stroke plane between the power stroke (PS) and recovery stroke (RS) to minimize appendage collision (Schabes and Hamner 1992). Species with lower *G/L*, such as copepods, isopods and stomatopods, typically perform a “hybrid metachronal stroke” consisting of a metachronal power stroke followed by a synchronous or near-synchronous recovery stroke (Alexander 1988; van Duren and Videler 2003; Kiørboe et al. 2010; Campos et al. 2012). In particular, copepods and stomatopods swim intermittently for escaping, feeding and other rapid maneuvers. Calanoid copepods (*Calanus finmarchicus*) reach 7.4 times the acceleration due to gravity during escape jumps (Murphy et al. 2012). Normalized speeds as fast as 40 body lengths/s have been reported during escape swimming of mantis shrimp using hybrid metachronal rowing (Campos et al. 2012). The range of *θ* used during escape swimming of mantis shrimp is larger than the range of *θ* used during hovering and fast forward swimming gaits of Antarctic krill (Murphy et al., 2011). Given that a comparative assessment of the mechanical performance of pure and hybrid metachronal rowing is currently unavailable, it is unknown whether the hybrid rowing strategy improves burst swimming.

For a given morphology (*G/L*) and paddle tip speed, we hypothesize that hybrid metachronal rowing can generate greater acceleration and swimming speed as compared to pure metachronal rowing by permitting operation at larger *θ* without the concern of appendage collisions. Although several studies have examined the wake generated by metachronal rowing across different species and behaviors (Yen et al. 2003; Lim and DeMont 2009; Catton et al. 2011; Murphy et al. 2013), the differences in body morphology and behaviors across species make it difficult to isolate the effects of kinematic parameters on swimming performance. We characterize the hybrid metachronal rowing kinematics of *Neogonodactylus bredini* mantis shrimp during burst swimming and compare these results to simulations of rigid paddles to ascertain whether hybrid metachronal rowing permits operation at larger *θ* as compared to pure metachronal rowing. We next use a self-propelling robot (Ford and Santhanakrishnan 2021a, 2021b) to compare swimming speed, peak acceleration, and wake structure of synchronous (*ϕ*=0%), pure metachronal (*ϕ*=10-20%), and hybrid metachronal (*ϕ*=10-20%).

## Methods

### Care and recording of live animals

Adult mantis shrimp (Crustacea: Stomatopoda: Gonodactylidae: *Neogonodactylus bredini*) were collected at the Galeta Marine Station, Smithsonian Tropical Research Institute, Panama (Permit # SC/A-6-19). Each mantis shrimp was housed in individual tanks within a recirculating saltwater system at Duke University (44 liter tanks, 12 h:12 h light:dark cycle; 27–28° C, salinity 32–36 parts per thousand). The mantis shrimp were fed a combination of live snails and defrosted krill three times per week. Over the course of 3 weeks, mantis shrimp were fed in the presence of increasing light intensity to acclimate them to the bright lights used during high-speed imaging.

Swimming trials were conducted using four animals in their individual home tanks and filmed using high-speed imaging (500 frames s^-1^; 1024×1024 pixel resolution; Fastcam SA-X2 and SA-Z, Photron, San Diego, CA, USA). Each tank was equipped with a white acrylic sheet background for contrast and angled in congruence with the light source (75 W LED, Varsa Nila, Inc., Altadena, CA, USA) for illumination. We filmed mantis shrimp swimming within the high-speed camera’s focal plane by inducing them to pursue bait held in a pair of forceps. A calibration ruler was positioned in this plane of focus to calibrate the images. Body length (BL) of the individuals ranged from 50.6 to 57.9 mm (**see Table S1** in **Supplementary Material** for details).

### Modeling appendage collisions

Stroke amplitude is constrained by the potential for appendage collision during pure metachronal rowing, given that maximum stroke amplitude is a function of *G/L* and phase lag. We calculated the tip-to-tip distance between two simulated rigid paddles with dimensionless length *L*=1 and dimensionless gap between paddles *G*=0.56. The *G* and *L* values were based on the lowest *G/L* value of 0.56 that was measured among the four *N. bredini* individuals. The tip-to-tip distances were evaluated in order to determine how varying phase lag in pure metachronal rowing changed the maximum stroke amplitude before the two paddles collide at some point in the cycle. We calculated the combinations of *θ* and *ϕ* that result in collisions between adjacent pleopods. Each paddle was oscillated using a simple harmonic motion profile given by the following relation:

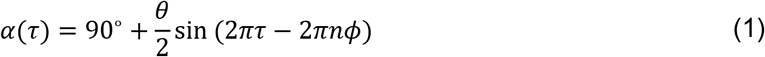

where *α*(*τ*) is the paddle angle at dimensionless time instant *τ* (**Figure 1A**). The non-dimensional stroke time *τ* runs from 0 (start of a cycle) to 1 (end of a cycle). *n*=0 for the posterior paddle and *n*=1 for the anterior paddle. Both paddles were oscillated with the same stroke amplitude *θ*, where *θ* was varied from 0° to 180° for the simulations. Phase lag *ϕ* was varied from 0% to 25% of cycle duration (*τ*=1). The horizontal distance between the tips of the oscillating rigid appendages (*d*) was calculated at each dimensionless time instant *τ* according to the following equation:

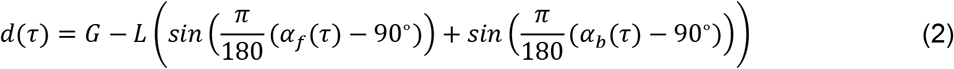

where *α_f_* is the angle of the forward/anterior paddle at time *τ*, and *α_b_* is the angle of the more backward/posterior paddle at time *τ. d* ≥ 0 corresponds to a position where the two paddles do not collide, and this was considered to be an “achievable” position. The paddles collide with each other when *d* < 0 and this was considered to be an “unachievable” position. Simulations were conducted for varying *ϕ* and *θ* in MATLAB (The MathWorks Inc, Natick, NJ, USA; version 9.9.0) in order to determine whether collisions occur at any point in a cycle (i.e., *d* < 0). Combinations of *ϕ* and *θ* that had achievable (i.e., *d* ≥ 0) positions at every instant within a stroke cycle were considered to be achievable kinematics for a pure metachronal stroke. We compared the simulation results to the stroke amplitude and phase lag values tracked from the recordings of *N. bredini* to determine whether the pleopod kinematics used by the mantis shrimp during hybrid metachronal rowing exceeded the maximum values achievable by pure metachronal rowing.

**Figure 1:**
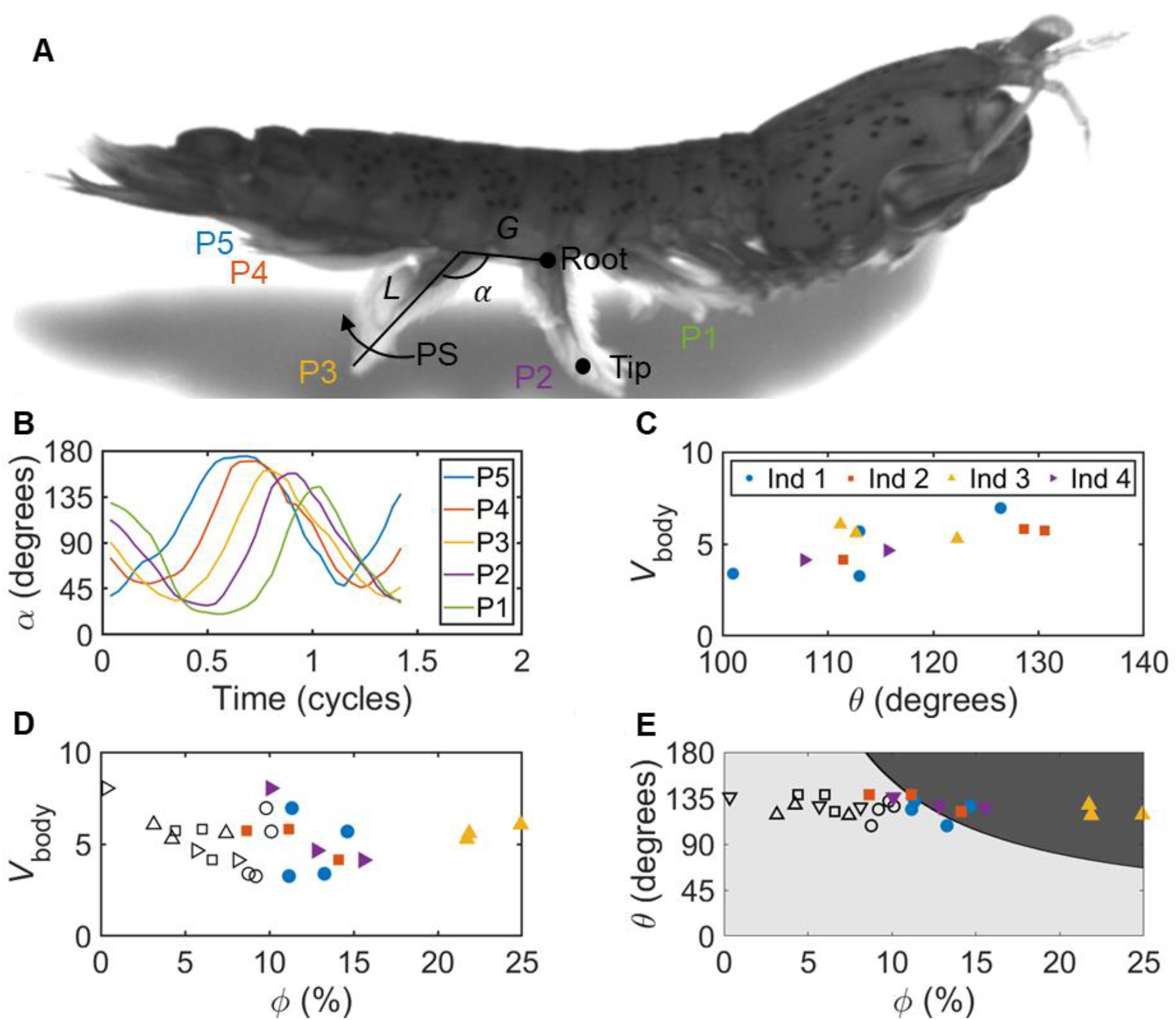
(A) Lateral view of an *N. bredini* individual (extracted from a high-speed video) indicating pleopods (P1-P5, where P1= anterior pleopod and P5= posterior pleopod), inter-pleopod spacing (*G*), pleopod length (*L*), pleopod angle (*α*) and the direction of the power stroke (PS). (B) Example of time-variation of pleopod angles of P1-P5 in non-dimensional time. The delay between the motion of adjacent pleopods is the phase lag, which is larger during power stroke than during recovery stroke. The amplitude of a pleopod angle waveform indicates the pleopod stroke amplitude (*θ_Pn_*). (C) Mean stroke amplitude (*θ*, average of stroke amplitudes of 5 pleopods) and swimming speed showed marginal variation between different individuals (N=4). Multiple markers for an individual indicate the outcomes of different trials conducted under identical test conditions. (D) Phase lag (*ϕ*) was generally lower in recovery stroke (hollow markers) as compared to power stroke (filled markers). (E) For pure metachronal rowing at a given *G/L*, stroke amplitude is limited by collision between the tips of adjacent appendages. Simulations of 2 rigid paddles undergoing simple harmonic motion (*G/L*=0.56 corresponding to *N. bredini*) show that achievable stroke amplitudes of pure metachronal rowing (shown in light gray region) is a function of phase lag. Combinations of *ϕ* and *θ* used by *N. bredini* (N=4) largely fall outside the light gray region achievable using pure metachronal rowing, showing that hybrid metachronal rowing can permit operation at larger stroke amplitudes without inter-appendage collisions.

Contour plots showing the achievable and unachievable kinematics for the mantis shrimp (*G/L*=0.56) and for the robotic model (*G/L*=0.5) are shown in **Figure 1** and **Figure 2**, respectively. The test conditions selected for use in this study are indicated in Figure 2E, where kinematics achievable for a pure metachronal stroke are indicated in light gray, while kinematics unachievable for a pure metachronal stroke are indicated in dark gray. The hybrid metachronal stroke allows for larger stroke amplitude for a constant phase lag and nondimensional ratio of gap between paddles to paddle length.

**Figure 2:**
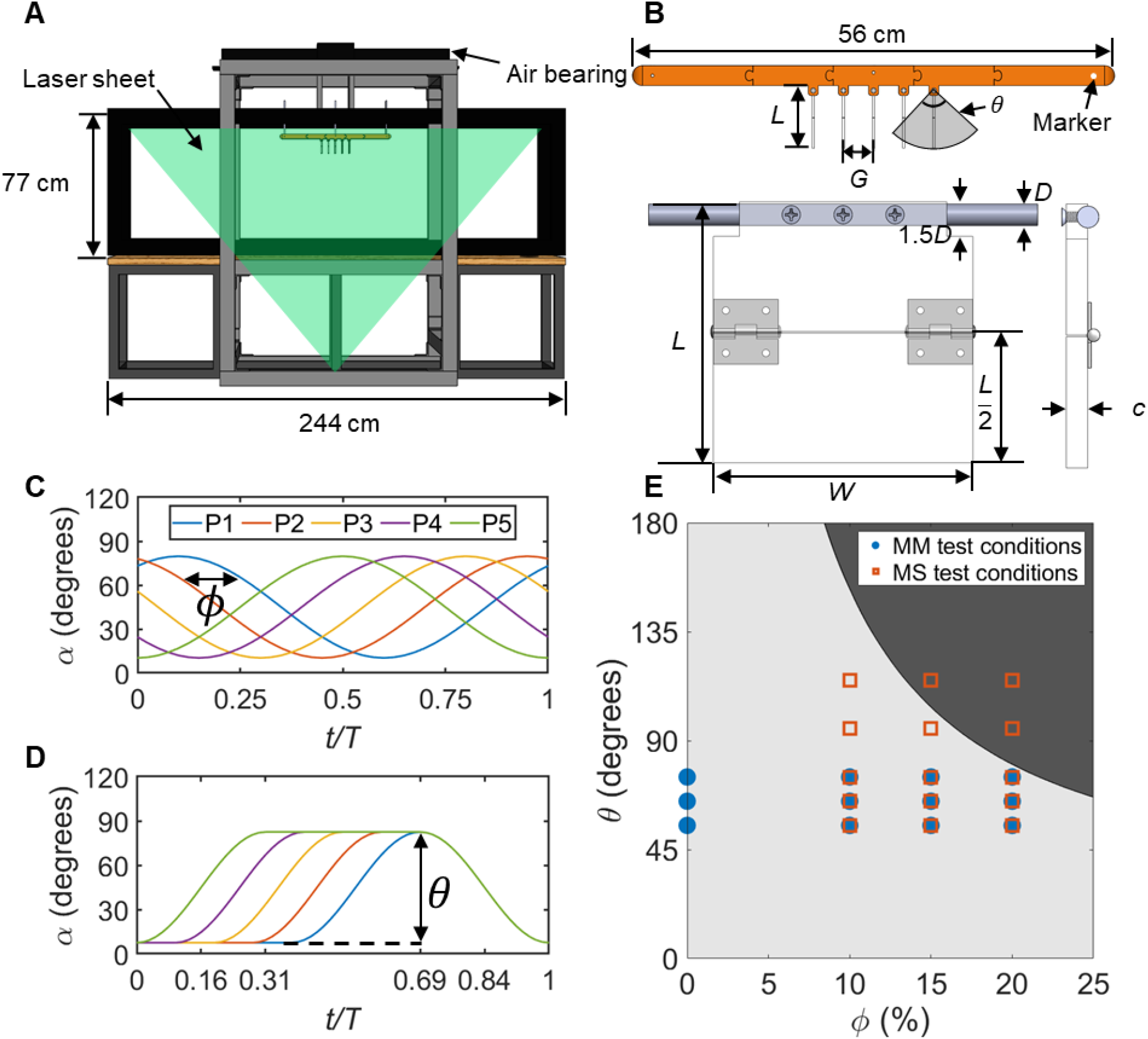
Experimental apparatus and test conditions for the robotic model used in this study. (A) Assembled model suspended from a linear air bearing (for self-propulsion) and submerged in a 2.44 m long aquarium, with laser sheet used for 2D time-resolved particle image velocimetry (PIV) measurements shown. (B) Schematic of the robotic model consisting of 5 flat-plate paddles and magnified view of an individual paddle. The paddle length (*L*=7.62 cm), paddle width (*W*=7.62 cm), and paddle thickness (*c*=0.32 cm) are shown. The PIV laser sheet cuts through the vertical mid-plane of the paddles. (C) Prescribed model kinematics (paddle angle *α*) for pure metachronal rowing, with time points 0%, 50%, and 100% of both power stroke and recovery stroke shown. Time (*t*) is non-dimensionalized by cycle time (*T*). Note that for a pure metachronal stroke, 0% power stroke corresponds to *t*/*T*=0, 50% power stroke corresponds to *t*/*T*=0.25, 100% power stroke and 0% recovery stroke correspond to *t*/*T*=0.5, 50% recovery stroke corresponds to *t*/*T*=0.75, and 100% recovery stroke corresponds to *t/T=1*. (D) Example of prescribed kinematics for hybrid metachronal rowing (*α* versus *t*/*T*) with 0%, 50%, and 100% of each power and recovery strokes labeled. For a hybrid metachronal stroke, the P5 paddle pauses after the power stroke, so the total cycle time (*T*) increases relative to pure metachronal rowing, and the *t*/*T* values at which the indicated time points occur to change and the 100% power stroke and 0% recovery stroke time points to become distinct from each other. (E) Test conditions used in this study overlaid on the achievable (light gray)/unachievable (dark gray) combinations of *θ* and *ϕ* obtained from modeling 2 rigid paddles (*G/L*=0.5) undergoing simple harmonic motion for pure metachronal rowing (MM). *θ* was varied from 55° to 75° for synchronous and pure metachronal (MM) rowing. *θ* was varied from 55° to 115° for hybrid metachronal rowing (MS). For each *θ* considered in pure metachronal (MM) and hybrid rowing (MS) kinematics, *ϕ* of 10%, 15% and 20% were tested, in addition to *ϕ*=0% corresponding to synchronous rowing.

### Robotic model

A self-propelling metachronal paddling robot was developed previously (Ford and Santhanakrishnan, 2021a, Ford and Santhanakrishnan, 2021b) to perform cross-species comparisons and identify the functional roles of morphological and kinematic parameters on metachronal swimming performance. We used this robot to compare the performance of hybrid and pure metachronal rowing kinematics. Motion of the paddles was driven by pulleys attached to five stepper motors. A high-speed camera (Phantom Miro M110, Vision Research, Wayne, NJ, USA) with maximum frame rate of 1688 frames/second at full resolution (1280 x 800 pixels) was placed 124 cm from the front wall of the aquarium with a 60 mm fixed focal length lens (aperture set to f/2.8). Recordings were acquired at 250 frames/s (100 frames per stroke cycle for pure metachronal rowing, 100(1+4*ϕ*) frames per cycle for hybrid metachronal rowing) with the robotic model performing different kinematics. Robot displacement was tracked from the video recordings, and the displacement data were post-processed to determine swimming speed and peak acceleration.

The body of the robot was 56 cm in length and used square paddles 7.62 cm in both width (*W*) and length (*L*), with an inter-paddle gap (*G*) of 3.81 cm between adjacent paddles. The gap and length values were selected to obtain *G/L*=0.5, which is within the biological range of 0.2≤*G*/*L*≤0.7 that was previously reported across a number of metachronal swimming species (Murphy et al. 2011). The width of the paddles was made equal to the length to be representative of the low aspect ratio (length divided by width) pleopods of mantis shrimp, relative to the high aspect ratio pleopods of copepods and euphausiids (Campos et al. 2012). The model was submerged in a solution of 85% glycerin (density=1220 kg/m^3^, kinematic viscosity=100 mm^2^/s), in a glass aquarium measuring 244 cm in length, 65 cm in width, and 77 cm in height (**Figure 2 A-B**). The model was suspended from a 1 m long air bearing which allowed for linear motion with minimal friction (**Figure 2A**), and which restricted the model position to be 31 cm from both of the side walls, 63 cm from the lower surface of the tank, 5 cm below the water surface, and at least 60 cm from the ends of the tank in the direction of motion.

Time-variation of paddle root angle (*α*) was prescribed for synchronous rowing, pure metachronal rowing and hybrid metachronal rowing (i.e., metachronal power stroke followed by a pause and synchronous recovery stroke), with stroke amplitude (*θ*) being varied from 55°-115°. Phase lag (*ϕ*) in both power and recovery strokes during pure metachronal rowing was varied from 10% to 20% of the complete cycle duration. *ϕ* in both power and recovery strokes was maintained at 0% of the cycle duration for synchronous rowing. For hybrid metachronal rowing, *ϕ* in power stroke was varied from 10% to 20% of the combined duration of the power and recovery strokes (excluding the pause). Also, for hybrid metachronal rowing, *ϕ* in recovery stroke was maintained at 0% of the cycle duration (same as the recovery stroke in synchronous rowing). For hybrid metachronal rowing, each paddle began its respective power stroke in sequence, and then paused following the completion of the power stroke until all paddles completed their motion. Once the anterior paddle had completed its power stroke, all five paddles performed their recovery strokes synchronously. This allowed the paddles to achieve the same maximum tip speeds for the same phase lag and stroke amplitude conditions, regardless of whether they were performing synchronous, pure metachronal, or hybrid metachronal rowing kinematics.

Stroke period for synchronous and pure metachronal rowing was *T*=0.4 seconds, which corresponds to stroke frequency *f* = 1/*T*=2.5 Hz. For hybrid metachronal rowing, the stroke period varied with *ϕ* due to pausing at the end of power stroke, so that *T*=0.4(1+4*ϕ*) seconds and stroke frequency (*f* = 1/*T*) ranged from 1.78 Hz for *ϕ*=10% to 1.39 Hz for *ϕ*=20%. Examples of metachronal and hybrid kinematics with *θ* = 75° and *ϕ* = 15% are shown in **Figure 2 C-D**, with time points representing 0%, 50%, and 100% of the P5 power stroke and P5 recovery stroke indicated. For a pure metachronal stroke, the end of power stroke and the start of recovery stroke coincide, while for a hybrid metachronal stroke of the P5 paddle the end of power stroke and the start of recovery stroke do not.

Reynolds number (*Re*) was based on the paddle length (*L*), stroke amplitude (*θ*), stroke period (*T*)), using the same equation as in Ford and Santhanakrishnan (2021b):

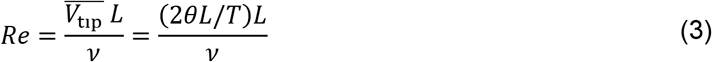

where 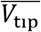 is the mean appendage tip speed 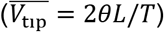 and the *v* is the kinematic viscosity of the fluid. As *θ* was varied, *Re* for robotic model tests was in the range of 155 to 416. This *Re* range was in the middle of the *Re*~10^1^-10^3^ range observed in intermittently swimming crustaceans, including copepods (Morris et al. 1990; Lenz et al 2004; Kiørboe et al. 2010) and stomatopods (Campos et al. 2012, present study).

As scaled-up paddles were used in the robot that was submerged in a large volume of water-glycerin mixture (100 times more viscous than water), and as *Re* is proportional to the square of appendage length, the robot was required to operate at stroke frequencies (*f* = 1/*T*) lower than those observed in *N. bredini* in order to achieve *Re* in the biological range. The lower stroke frequencies used by the robotic model when compared to those of free-swimming stomatopods (Campos et al. 2012; present study) are expected to result in the robot having lower swimming speed. However, as the maximum paddle tip speed for a given stroke amplitude was maintained equal between pure and hybrid metachronal rowing, we expect comparisons of the swimming speed and peak acceleration to be unaffected by our approximation of using a lower stroke frequency.

### Swimming performance

#### Organismal kinematics

From the recordings of live mantis shrimp *N. bredini*, we calculated phase lag, *ϕ*, for both the power stroke and recovery stroke and swimming speed (*V*_body_). Measurements of swimming speed and pleopod root angle (*α*) were performed in ImageJ software (Schneider et al. 2012; version 1.52a). A representative image of an individual is shown in **Figure 1A**, indicating the five pleopods numbered sequentially from anterior (P1) to posterior (P5), gap between pleopods (*G*) and pleopod length (*L*). *α*, defined relative to the longitudinal body axis, was determined for a minimum of 20 times per cycle (example of tracked α is shown in **Figure 1B**). While *θ* of each pleopod is similar, the phase lag *ϕ* (temporal separation between α profiles of adjacent pleopods) is much larger during power stroke than during recovery stroke. There is often a pause between the power and recovery strokes which results in multiple time points at which the pleopods are at their maximum angle, or an incomplete pause in which the pleopod angular motion slows but does not completely stop, which can result in the time at which the pleopod reaches maximum *α* being delayed for some pleopods but not others. Because there is often a complete or incomplete pause, it can be misleading to determine *ϕ* only by measuring the time delay between adjacent pleopods achieving their maximum or minimum root angle (*α*_max_ or *α*_min_). For a more robust characterization of *ϕ*, we defined *ϕ* as the time between each pleopod reaching peak angular velocities during both the power and recovery strokes, divided by the stroke period (duration of power stroke duration of recovery stroke). This definition offers the advantage of allowing for calculation of *ϕ* in the power stroke and recovery stroke independently. In addition to tracking pleopod kinematics, we tracked the position of these organisms throughout the swimming cycle using ImageJ software in order to determine their average swimming speed.

#### Robot swimming performance

The robotic model was recorded swimming using a variety of paddle kinematics as described earlier (under the subheading Robotic Model). Position of the robot was tracked in time using DLT program (Hedrick 2008, version 7), and a custom MATLAB code was used to determine the velocity and acceleration of the model in time. Average velocity and acceleration values were determined over the period of one cycle (0.4 seconds for synchronous and pure metachronal rowing, 0.4·(1+4*ϕ*) seconds for hybrid metachronal rowing).

### Particle image velocimetry (PIV)

Two-dimensional time-resolved PIV measurements were used to visualize the flow generated by paddling under varying kinematic conditions. 55-micron diameter silver-coated polyamide particles (LaVision GmbH, Göttingen, Germany) were used for seeding the flow for PIV recordings. The same high-speed camera used to record swimming performance (Phantom Miro M110, Vision Research, Wayne, NJ, USA) was placed 110 cm from the front of the aquarium with the lens and aperture settings as used for swimming speed measurements (see the subheading Robotic model). Illumination was provided using a single-cavity high-speed Nd:YLF laser with 527 nm wavelength, capable of providing maximum 30 mJ/pulse at a pulse rate of 10 kHz (model DM-527, Photonics Industries International Inc., Ronkonkoma, NY, USA). Recordings were performed at the same frame rates that were used for displacement tracking so as to obtain 100 velocity vector fields per paddling cycle. Multi-pass PIV cross-correlation was performed in DaVis 8.4 software (LaVision GmbH, Göttingen, Germany) with one pass of 48×48 pixels with 50% overlap, and two passes of 24×24 pixels with 50% overlap for each pass. Vector post-processing was performed for the final pass to eliminate spurious vectors with Q>1.3 (where Q is the ratio of highest correlation coefficient divided by second highest correlation coefficient) and empty spaces were re-filled with interpolated values.

### Calculated Quantities

The swimming speed of mantis shrimp individuals was calculated from position data tracked from each recording according to the following equation:

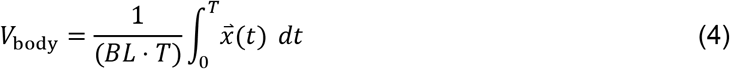

where *V*_body_ is the swimming speed of the organism, *T* is the duration of the paddling cycle period tracked from the high-speed recordings, which ranged from 0.1 seconds to 0.15 seconds, *t* = 0 seconds is the start of a power stroke for the fifth pleopod, 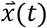 is the position of the organism at time *t* and *BL* is the body length of the organism.

Average *ϕ* between the motion of adjacent pleopods was calculated separately for power stroke and for recovery stroke in each recording. Since there was often a pause between the end power stroke and the start of recovery stroke (this pause being the defining characteristic of hybrid metachronal rowing), *ϕ* could not be calculated based solely on the maximum and minimum values of the pleopod root angle (*α*, **Figure 1**), since there could be multiple local extrema or the pleopod motion could slow without completely stopping, confounding the results. *ϕ* was therefore defined based on the duration of time between when adjacent pleopods reached their maximum angular velocities. In order to minimize uncertainty, the overall P5-P1 phase lag was calculated, and the average was used to define *ϕ*:

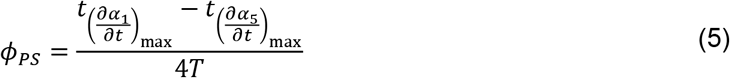

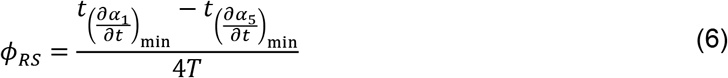

where *ϕ_PS_* and *ϕ_RS_* are the average phase lags during the power stroke and recovery stroke, respectively. 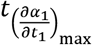 and 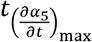 are the times at which the first and fifth pleopods, respectively, reach their maximum angular velocities during power stroke, while 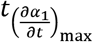 and 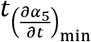 are the times at which the first and fifth pleopods reach their maximum angular velocities during the recovery stroke. The factor of 1/4 is introduced to the equation in order to determine the average phase lag (because there are 5 pleopod pairs, there are 4 phase lags).

2D velocity vector fields obtained from PIV measurements, containing horizontal (*x*-axis) component of velocity (*u*) and vertical (*y*-axis) component of velocity (*v*) within the PIV field of view, were used to visualize the wake generated by paddling. Vorticity is a measure of angular rotation in a fluid flow and has been linked to the generation of propulsive forces (Wu 1980). Out-of-plane component of vorticity (*ω_z_*) was calculated from the 2D velocity vector fields according to the equation:

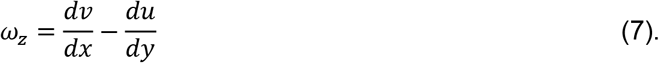

Momentum flux is a surrogate measure of the force imparted to the fluid by the paddling motion (Ford et al. 2019). The rate of transfer of momentum to the fluid is directly related to the generation of propulsive lift and drag forces necessary to maneuver underwater. Cycle-averaged momentum fluxes in the horizontal and vertical directions were calculated at several locations for each kinematic condition tested in this study. Horizontal momentum flux 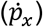 and vertical momentum flux 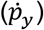 were defined according to the following equations:

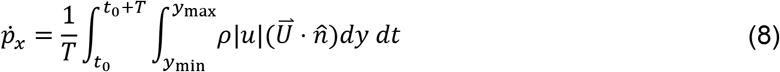

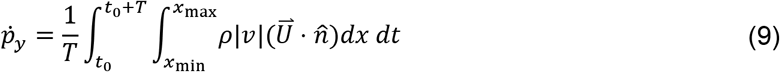

where *t_0_* is the time at the start of a paddling cycle, *ρ* is the density of the fluid, 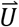 is the velocity vector at a given location in the flow field, and 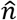 is the unit vector in the direction of interest (horizontal for horizontal momentum flux and vertical for vertical momentum flux).

Robot swimming speed was calculated from the linear displacement averaged over one stroke period, as defined in the equation below:

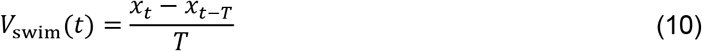

where *V*_swim_(*t*) is the average swimming speed over a cycle at time *t, x_t_* is the position of the robot at time *t, x_t-T_* is the displacement of the robot one stroke period (*T*) before time *t*. The cycle-averaged velocity over the last cycle was defined as *V*_swim_ and is presented in **Figure 5A**.

Acceleration from rest is expected to be an important measure of burst swimming performance, rather than other common measures of efficiency used for steady swimming performance (Walker 2002). We calculated time-varying acceleration of the robotic model according to the equation below:

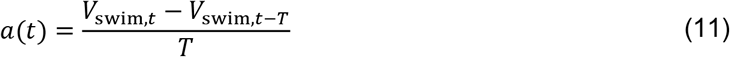

where *a*(*t*) is the acceleration of the robot at time *t*. The maximum value of the acceleration was calculated and is presented in **Figure 5B**.

## Results

### Organismal swimming

We characterized the pleopod kinematics and swimming speed in 13 recordings acquired in 4 *N. bredini* individuals. The angles of the pleopods relative to the body axis (*α*) were tracked in time and a sample of the resulting profiles for one recording is shown in **Figure 1B**. A clear phase delay (*ϕ*) between different pleopods was observed during power stroke that was nearly constant along the body. Comparatively, a much smaller phase lag was observed during recovery stroke for each pleopod. This means that while all pleopods stroked metachronally during power stroke, the recovery stroke was near-synchronous. This is similar to the qualitative observations of kinematics reported in another stomatopod species (*O. havanensis*, Campos et al., 2012).

Based on the simulation of pleopod collisions using pure metachronal rowing, we found that several of the phase lags recorded during the power strokes of the mantis shrimp fall into the unachievable kinematics region, but all of the phase lags observed in the recovery strokes fall into the achievable region. This could indicate that mantis shrimp selectively use the hybrid metachronal stroke to maximize *θ* and achieve large forward swimming speeds. Based on these observations, we used a self-propelling robot that performed synchronous, pure metachronal, and hybrid metachronal kinematics to answer whether hybrid metachronal kinematics provides performance benefits relative to pure metachronal kinematics at the same stroke amplitude and maximum tip speed, and how increasing *θ* affected body acceleration.

### Flow generated by pure and hybrid metachronal rowing

Wake flow fields were extracted at the start, middle and end of power stroke, and at the start, middle and end of recovery stroke (time instants based on the posterior paddle). For a fixed stroke amplitude, the pure metachronal wake shows a more continuous jet compared to the periodic wakes generated by paddling with synchronous and hybrid metachronal kinematics. Additionally, the vorticity is stronger away from the body in pure metachronal rowing than in synchronous or hybrid metachronal rowing, with the wake directed in a more vertical direction. Counter-rotating vortices generated by the paddles during the power and recovery strokes interact constructively during pure metachronal rowing, which helps direct the flow into a jet. These constructive interactions are not present in either synchronous or hybrid metachronal rowing. For hybrid metachronal rowing with *θ* = 115°, there is visible flow directed in the anterior direction which is not seen under any of the three types of kinematics when *θ* = 75°.

In order to quantify the downstream wake generated by the paddling robot, we calculated momentum flux in both the vertical and horizontal directions at select locations in the flow (**Figure 4**). In general, *ϕ* was found to have minimal effect on either the horizontal 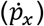 or vertical 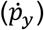 momentum flux when compared to the stroke amplitude. The mean values of momentum flux for pure metachronal rowing motion (particularly in the vertical direction, 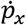) were slightly higher than for hybrid metachronal rowing motion for the same location, same *ϕ*, and same *θ*, but the difference was relatively small and sometimes within the standard deviation based on the cycle-to-cycle variation of momentum flux in the wake. *θ* showed the strongest effect on the momentum flux in both the horizontal and vertical directions, and the variation between cycles increased with increasing *θ*. These results suggest that for constant stroke frequency, the stroke amplitude is the most important factor in determining the swimming speed. We next calculated the speed and acceleration of the robot from recordings of swimming with different prescribed kinematics.

**Figure 3:**
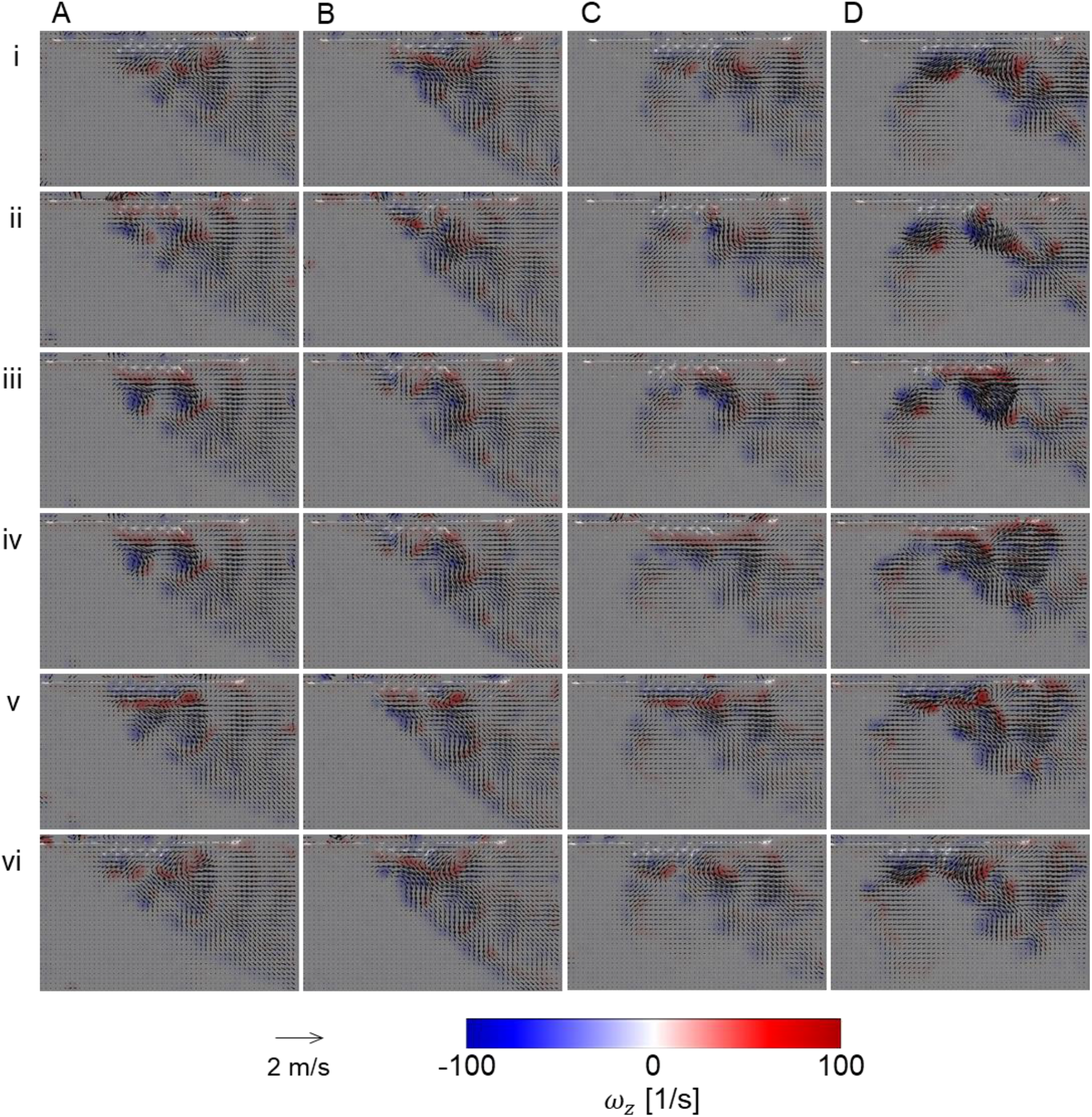
PIV velocity fields overlaid on contours of out-of-plane component of vorticity (*ω_z_*) for: (A) synchronous rowing, (B) pure metachronal rowing, (C) and (D) hybrid metachronal rowing. *ϕ* = 0% and *θ* = 75° in (A). *ϕ* = 15% and *θ* = 75° (B) and (C). *ϕ* = 15% and *θ* = 115° in (D). Time points shown include percentage of the power stroke (0%, row i; 50%, row ii; 100%, row iii) and recovery stroke (0%, row iv; 50%, row v; 100%, row vi). These time points correspond to *t*/*T*=0, 0.25, 0.5, 0.5, 0.75 and 1.0 for synchronous and pure metachronal kinematics, and *t*/*T*=0, 0.16, 0.31, 0.69, 0.84 and 1.0 for hybrid metachronal kinematics. Pure metachronal rowing result in the wake with the most clearly defined jet and also has the most vertical orientation. Hybrid metachronal rowing generates a more horizontally oriented wake. Increasing *θ* (as in D) results in a stronger wake with larger velocity, albeit with noticeable flow reversal near the end of power stroke (see row iii in column D).

**Figure 4:**
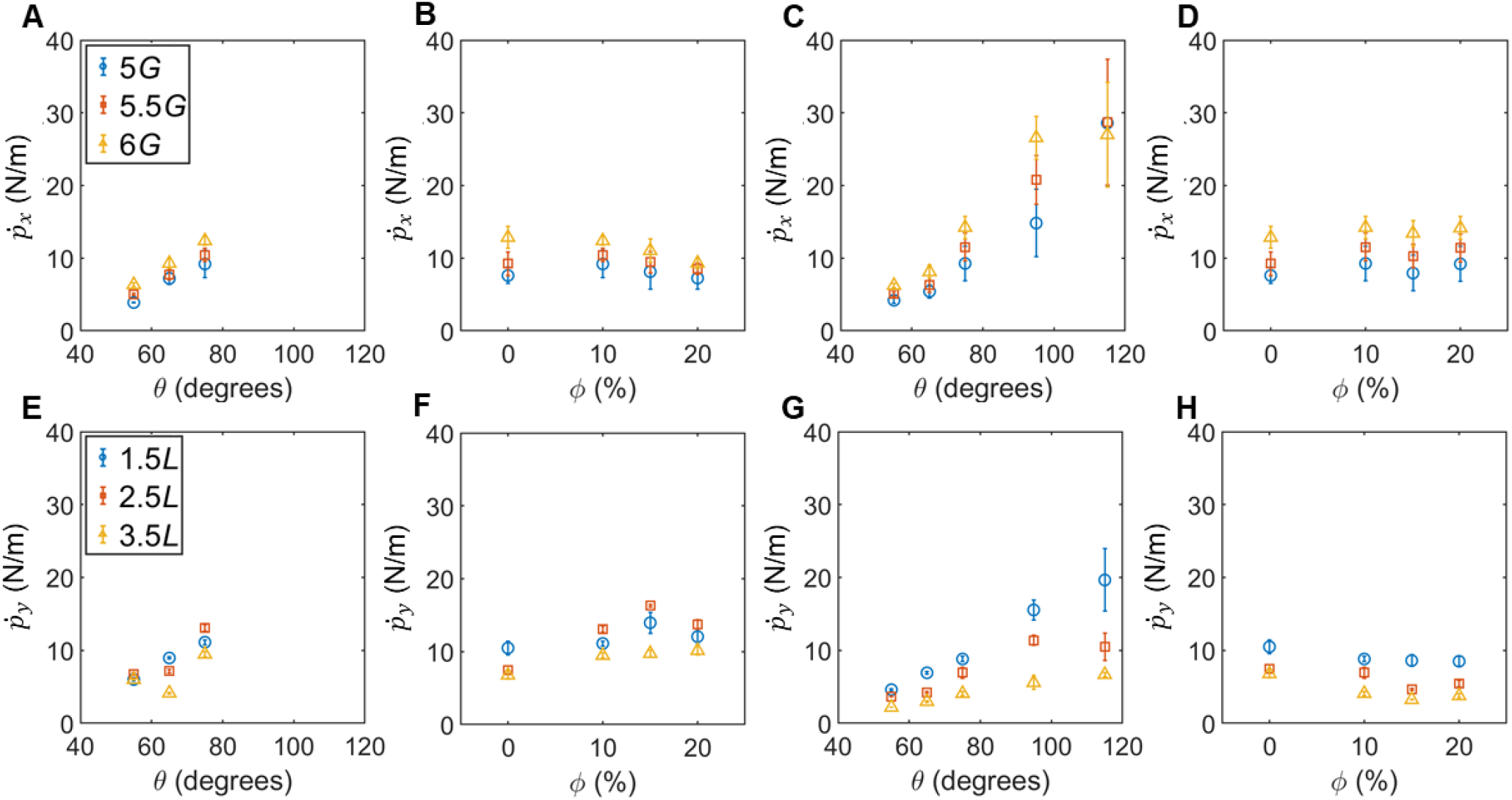
Cycle-averaged momentum fluxes as a function of stroke amplitude (*θ*) and phase lag (*ϕ*). (A)-(D) Horizontal momentum flux 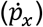 calculated at 5 to 6 inter-paddle gap (*G*) distances referenced from the anterior-most paddle, and vertical momentum flux 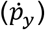 calculated at distances from 1.5 to 3.5 times of paddle length (*L*) referenced from paddle roots when at their most vertical position. 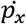 and 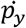 were calculated from cycle-averaged PIV velocity vector fields using equations (8) and (9), respectively. (A) 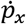 for pure metachronal rowing with *ϕ*=10%. (B) 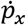 for synchronous (*ϕ*=0%) rowing and for pure metachronal rowing with *θ*=75°. (C) 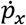 for hybrid metachronal rowing with *ϕ*=10%. (D) 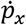 for synchronous and hybrid metachronal rowing with *θ*=75°. (E) 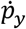 for pure metachronal rowing with *ϕ*=10%. (F) 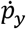) for synchronous and pure metachronal rowing with *θ*=75°. (G) 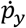 for hybrid metachronal rowing with *ϕ*=75°. (H) 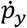 for synchronous and hybrid metachronal rowing with *θ*=75°. *ϕ* has a smaller effect on both 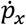 and 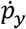 for hybrid metachronal kinematics (D, H) than *θ* (C, G). Pure metachronal kinematics has a larger maximum 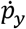 than 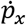 (vertically directed wake), while hybrid metachronal kinematics has a larger maximum 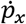 than 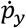 (horizontally directed wake).

### Swimming performance

The robot was allowed to swim along the air bearing according to the forces generated by the motion of the paddles. Position was tracked in time for the robotic model performing each set of prescribed kinematics with the cycle-averaged velocity and cycle-averaged acceleration calculated from the derivatives of the position data. Examples of the raw position, velocity, and acceleration data are shown in **Figure S1** in the **Supplementary Material**. Position data starts at 0 seconds while velocity and acceleration do not because velocity and acceleration were calculated via numerical differentiation of position across one cycle. The velocity at the end of travel (just before the robot reached the end of the air bearing) was determined for each condition and is shown in **Figure 5A**. For hybrid metachronal rowing, *ϕ* = 10% had the highest swimming speed for all *θ*, followed by *ϕ* = 15% and *ϕ* = 20%. However, for pure metachronal and synchronous rowing, the phase lag that results in the greatest swimming speed depends on the stroke amplitude. *θ* is an important factor in determining the swimming speed, but the influence of *θ* on swimming speed (i.e., δ*V_swim_*/δ*θ*) decreases with increasing *θ*. This could potentially be due to the forward-directed portion of the wake observed in the PIV results in **Figure 3**, which appears only for large *θ*. The advance ratio is a common measure of swimming performance in locomotor flows and is defined as the ratio of swimming speed to mean appendage tip speed (Walker 2002; Murphy et al. 2011). The robot did not always reach a steady swimming speed, and so the advance ratio was not calculated. It is also important to note that acceleration is far more important than advance ratio for burst swimming animals (such as *N. bredini*) that engage in rapid maneuvers starting from rest (when advance ratio equals zero).

**Figure 5:**
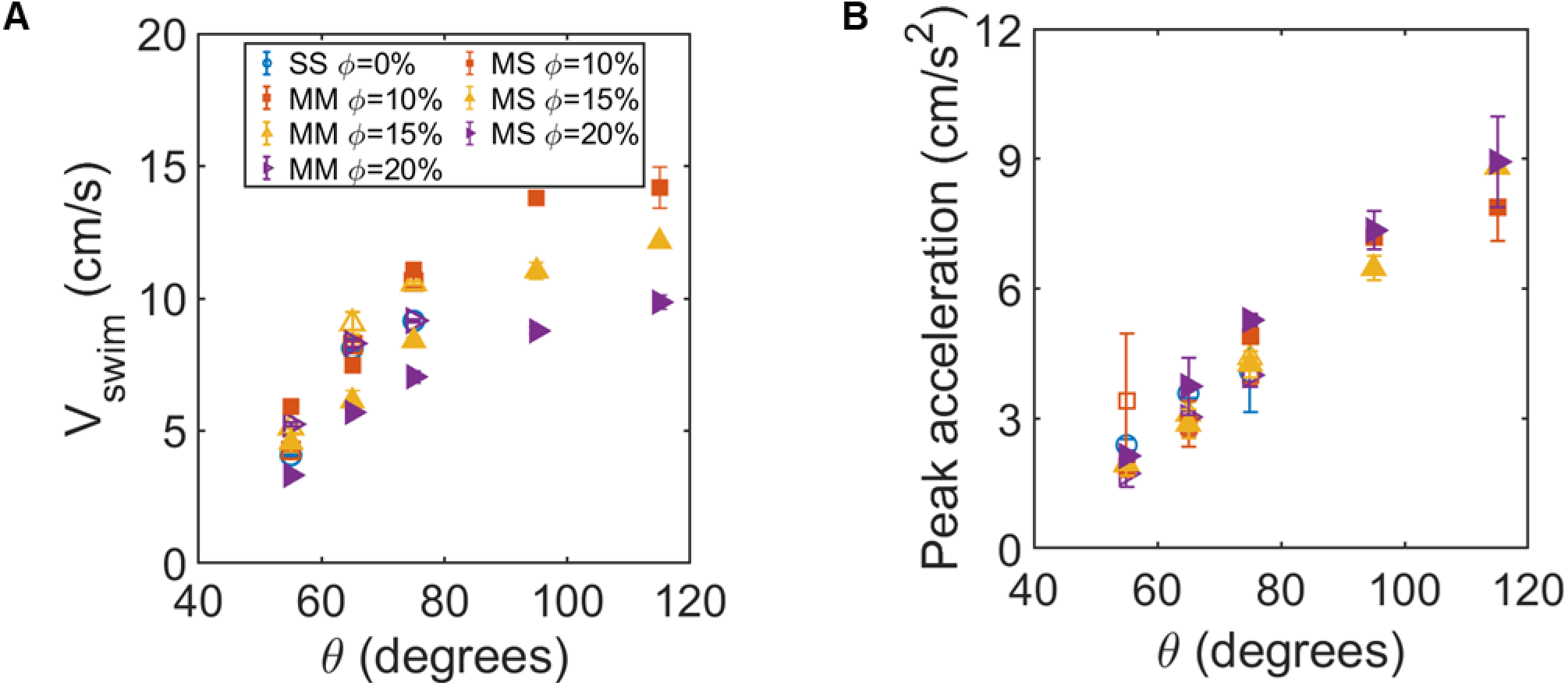
Swimming speed (*V*_swim_) and peak acceleration for the robotic model performing the various kinematics used in this study. (A) Swimming speed (*V*_swim_) increases with increasing stroke amplitude (*θ*) regardless of the type of kinematics, but the rate of increase of *V*_swim_ decreases with increasing *θ*. (B) Peak acceleration increased with *θ* for all conditions, but the rate of increase of peak acceleration did not decrease with increasing *θ*. SS represents synchronous rowing (hollow markers), MM represents pure metachronal rowing (hollow markers), and MS represents hybrid metachronal rowing (filled markers).

In addition to the velocity of the robot, we also calculated its peak acceleration. Peak acceleration is an important value in the locomotory performance of a burst swimming organism since rapid acceleration is necessary to perform the agile maneuvers to capture prey and/or avoid predation. The peak acceleration typically occurred after the first few paddling cycles had been completed and varied based on the phase lag and stroke amplitude of the kinematics being performed (see **Figure S1** in the **Supplementary Material**). While no clear trend in the peak acceleration could be determined based on changing the phase lag, stroke amplitude had a positive effect on the peak acceleration. Unlike the effect of stroke amplitude on velocity, which decreased with increasing *θ*, the effect of *θ* on peak acceleration did not decrease with increasing *θ*. This suggests that the large stroke amplitude achieved by using the hybrid metachronal kinematics is particularly well suited for acceleration from rest, rather than for sustaining high forward swimming speeds.

## Discussion

Using high-speed video recordings of mantis shrimp (*N. bredini*) individuals and experiments on a self-propelling robot, we tested whether hybrid metachronal rowing offers advantages for burst swimming as compared to pure metachronal rowing. The main findings of this study are: 1) stroke amplitude is generally a stronger predictor of metachronal swimming speed than phase lag for either pure metachronal or hybrid metachronal kinematics; 2) Hybrid metachronal kinematics can be used to surpass the stroke amplitude limitation of pure metachronal rowing, as the lower phase lag during the recovery stroke of hybrid metachronal kinematics minimizes opportunities for appendages to collide within a stroke cycle; and 3) larger peak acceleration and swimming speed were realized by the robot for hybrid metachronal kinematics when operating at a stroke amplitude that is in the range observed in *N. bredini* individuals than were achieved during pure metachronal rowing. Collectively, our findings suggest that intermittent swimmers with closely spaced rowing appendages may prefer hybrid metachronal kinematics over pure metachronal kinematics, as the phase lag during the recovery stroke was found to be smaller than the phase lag during the power stroke in every recording of swimming mantis shrimp. The likely reason for the preference of hybrid metachronal rowing is that it allows them to increase stroke amplitude and generate large accelerations needed for rapid maneuvering. While a previous study by Campos et al. (2012) qualitatively observed that the recovery stroke was nearly synchronous in another species of mantis shrimp (*O. havanensis*), the pleopod angles and phase lags were not characterized over a cycle as in this study.

Our use of a self-propelling robot allowed for the first-time comparison of mechanical performance of different types of stroke kinematics patterns (synchronous, pure metachronal, and hybrid metachronal) under identical test conditions. We used a stroke frequency that was lower than those of the *N. bredini* individuals (see **Table S1** in **Supplementary Material**), which resulted in lowered normalized swimming speeds as compared to *N. bredini* individuals. As we examined mechanical performance of synchronous, pure metachronal, and hybrid metachronal rowing under identical conditions, relative differences in swimming performance across the test conditions are expected to be unaffected by the reduced stroke frequency. Further experimentation is required to discern the role of stroke frequency in augmenting swimming speed. The use of both higher stroke frequencies and higher stroke amplitudes could explain the fast normalized swimming speeds observed in *O. havenensis* by Campos et al. (2012) when compared to *N. bredini* and to our robotic model.

Pure metachronal kinematics typically resulted in slightly higher swimming speeds than hybrid metachronal kinematics for the same stroke amplitude. However, varying the stroke amplitude had the largest effect on both velocity and peak acceleration of the robot. Increasing stroke amplitude augmented the swimming speed of the robot, but the rate of increase of swimming speed became smaller with increasing stroke amplitude. Peak acceleration of the robot also increased with increasing *θ*, but the rate of increase stayed nearly the same with increasing *θ*. The structure of the wake can help to explain why the effect of stroke amplitude on swimming speed decreases with increasing stroke amplitude. There is a region of reversed flow generated by the paddles with hybrid kinematics for *θ* = 115°. This flow will be detrimental to the swimming speed, since force generated by the paddling motion is directly related to the momentum transfer to the fluid. Additionally, the more structured wake of the pure metachronal kinematics results in a more vertically oriented jet than the hybrid metachronal kinematics. Though not the focus of this study, it is interesting to note that the more vertically oriented jets that originate from pure metachronal kinematics similar to those used by Antarctic krill (Murphy et al. 2011) could be conducive to hydrodynamic signaling between individuals when schooling (Catton et al. 2011; Murphy et al. 2019).

Several crustaceans, including copepods (van Duren and Videler 2003; Kiørboe et al. 2010), isopods (Alexander 1988) and stomatopods (Campos et al. 2012, this study), have been reported to use hybrid metachronal rowing. Unifying factors between these species are: 1) the use of appendage rowing for intermittent or burst swimming, including the use of rapid maneuvers for escaping and/or feeding; and 2) small inter-appendage spacing relative to appendage length (i.e., *G/L* ratio) that constrains use of large stroke amplitudes if using pure metachronal rowing. For organisms with long, closely spaced appendages such as copepods, flexibility can help to prevent damage from collisions between appendages. However, flexibility cannot increase maximum possible stroke amplitude when being constrained with a small *G/L* ratio. Hybrid metachronal rowing can be used to overcome this limitation and use a large stroke amplitude in order to achieve maximum acceleration for burst swimming behaviors.

The relative absence of hybrid metachronal rowing among continuously swimming organisms such as euphausiids (Murphy et al., 2011) suggests that the hybrid metachrony may not be as energetically efficient as pure metachrony. This is consistent with the fluid dynamic understanding that for Reynolds numbers much greater than 1, thrust is proportional to the square of the appendage tip velocity (i.e., *θ*^2^), while the appendage power input is proportional the appendage tip velocity cubed (i.e.,*θ*^3^; Blake 1979). Propulsive efficiency is defined as the ratio of swimming power divided by appendage power input and is therefore inversely proportional to the stroke amplitude. Hybrid metachronal rowing with large *θ* can provide greater peak acceleration at the expense of larger power input requirements and lower propulsive efficiency than pure metachronal rowing. While mechanical efficiency may need to be maximized for high endurance behaviors such as continuous or steady swimming, thrust generated in a stroke needs to be maximized for maneuvering behaviors (Walker 2002).

Metachronal rowing as a locomotion strategy is seen across a wide range of invertebrate sizes and shapes from diverse taxa (Lim and DeMont 2009; Kiørboe et al. 2010; Murphy et al. 2011; Campos et al. 2012; Funfak et al. 2015; Heimbichner Goebel et al. 2020). The benthic habitat of mantis shrimp versus pelagic habitat of euphausiids may play a role in shaping the appendage morphologies and stroke kinematics so as to meet specific locomotory needs. Hybrid metachronal rowing serves as an interesting example of how a locomotion strategy can be associated with periodic needs for high acceleration performance, as in copepods and stomatopods (Campos et al. 2012; Kiørboe et al. 2010). Our study indicates that by permitting large stroke amplitudes despite small inter-appendage spacing, hybrid metachronal rowing strategy is particularly well suited for the high acceleration requirements placed on burst swimming organisms such as *N. bredini*.

## Acknowledgements

The authors would like to thank Tyler Blackshare (Oklahoma State University) for assistance with the design and manufacturing of parts, C. Tanner Price (Oklahoma State University) for assistance in acquiring high-speed videos of robot motion, and Sophie Hanson (Duke University) for discussions and insights.

## FIGURES

**Table S1.**
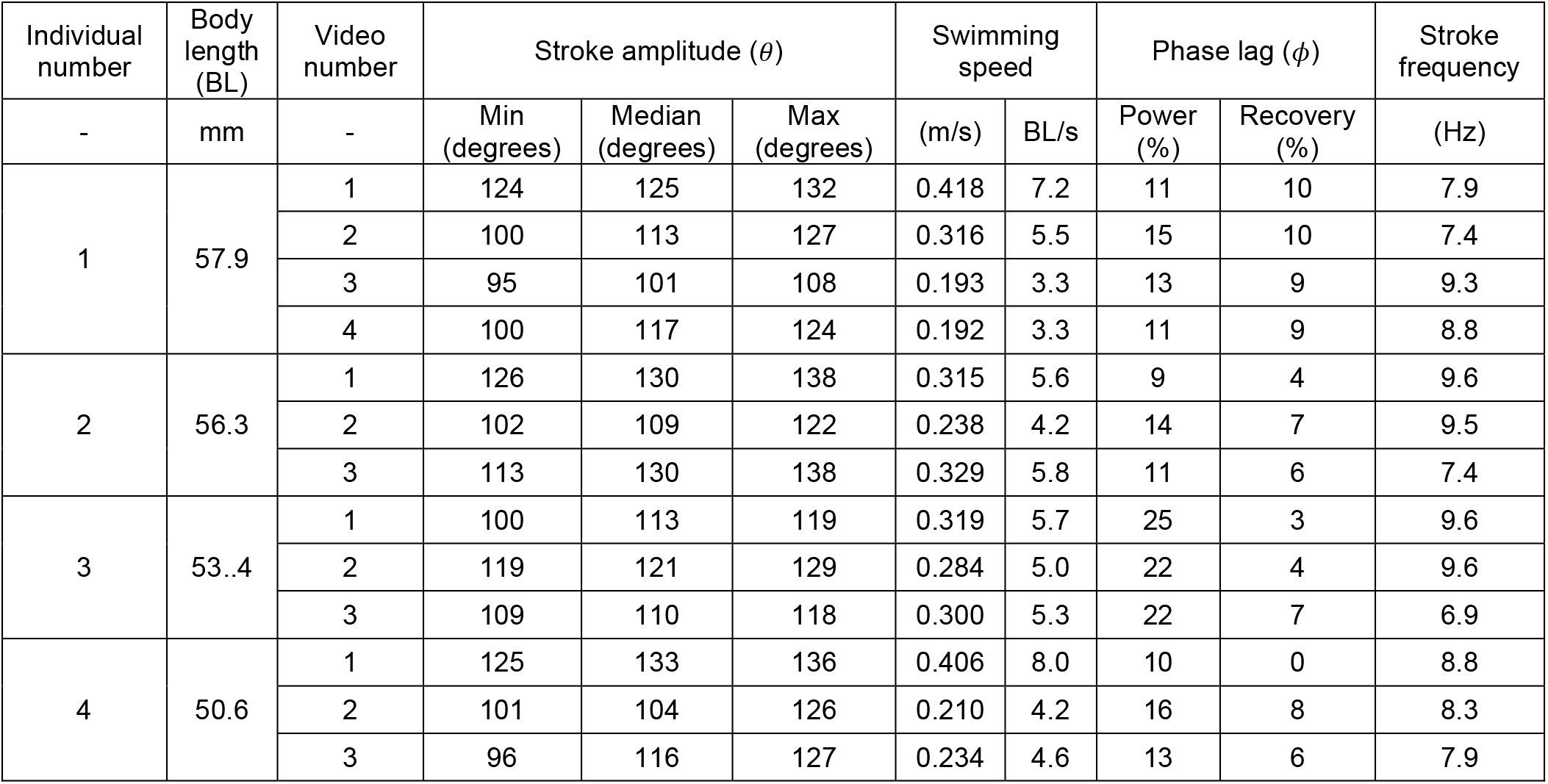
Swimming performance and kinematics data tracked from the recordings of *Neogonodactylus bredini* mantis shrimp (N=4) swimming used in this study. Swimming speed, mean stroke amplitude and mean phase lag are shown in **Figure 1**.

| Individual number | Body length (BL) | Video number | Stroke amplitude ($\theta$) | | | Swimming speed | | Phase lag ($\phi$) | | Stroke frequency |
| --- | --- | --- | --- | --- | --- | --- | --- | --- | --- | --- |
|  |  |  | Min (degrees) | Median (degrees) | Max (degrees) | (m/s) | BL/s | Power (%) | Recovery (%) |  |
| - | mm | - |  |  |  |  |  |  |  | (Hz) |
| 1 | 57.9 | 1 | 124 | 125 | 132 | 0.418 | 7.2 | 11 | 10 | 7.9 |
|  |  | 2 | 100 | 113 | 127 | 0.316 | 5.5 | 15 | 10 | 7.4 |
|  |  | 3 | 95 | 101 | 108 | 0.193 | 3.3 | 13 | 9 | 9.3 |
|  |  | 4 | 100 | 117 | 124 | 0.192 | 3.3 | 11 | 9 | 8.8 |
| 2 | 56.3 | 1 | 126 | 130 | 138 | 0.315 | 5.6 | 9 | 4 | 9.6 |
|  |  | 2 | 102 | 109 | 122 | 0.238 | 4.2 | 14 | 7 | 9.5 |
|  |  | 3 | 113 | 130 | 138 | 0.329 | 5.8 | 11 | 6 | 7.4 |
| 3 | 53..4 | 1 | 100 | 113 | 119 | 0.319 | 5.7 | 25 | 3 | 9.6 |
|  |  | 2 | 119 | 121 | 129 | 0.284 | 5.0 | 22 | 4 | 9.6 |
|  |  | 3 | 109 | 110 | 118 | 0.300 | 5.3 | 22 | 7 | 6.9 |
| 4 | 50.6 | 1 | 125 | 133 | 136 | 0.406 | 8.0 | 10 | 0 | 8.8 |
|  |  | 2 | 101 | 104 | 126 | 0.210 | 4.2 | 16 | 8 | 8.3 |
|  |  | 3 | 96 | 116 | 127 | 0.234 | 4.6 | 13 | 6 | 7.9 |

**Figure S1.**
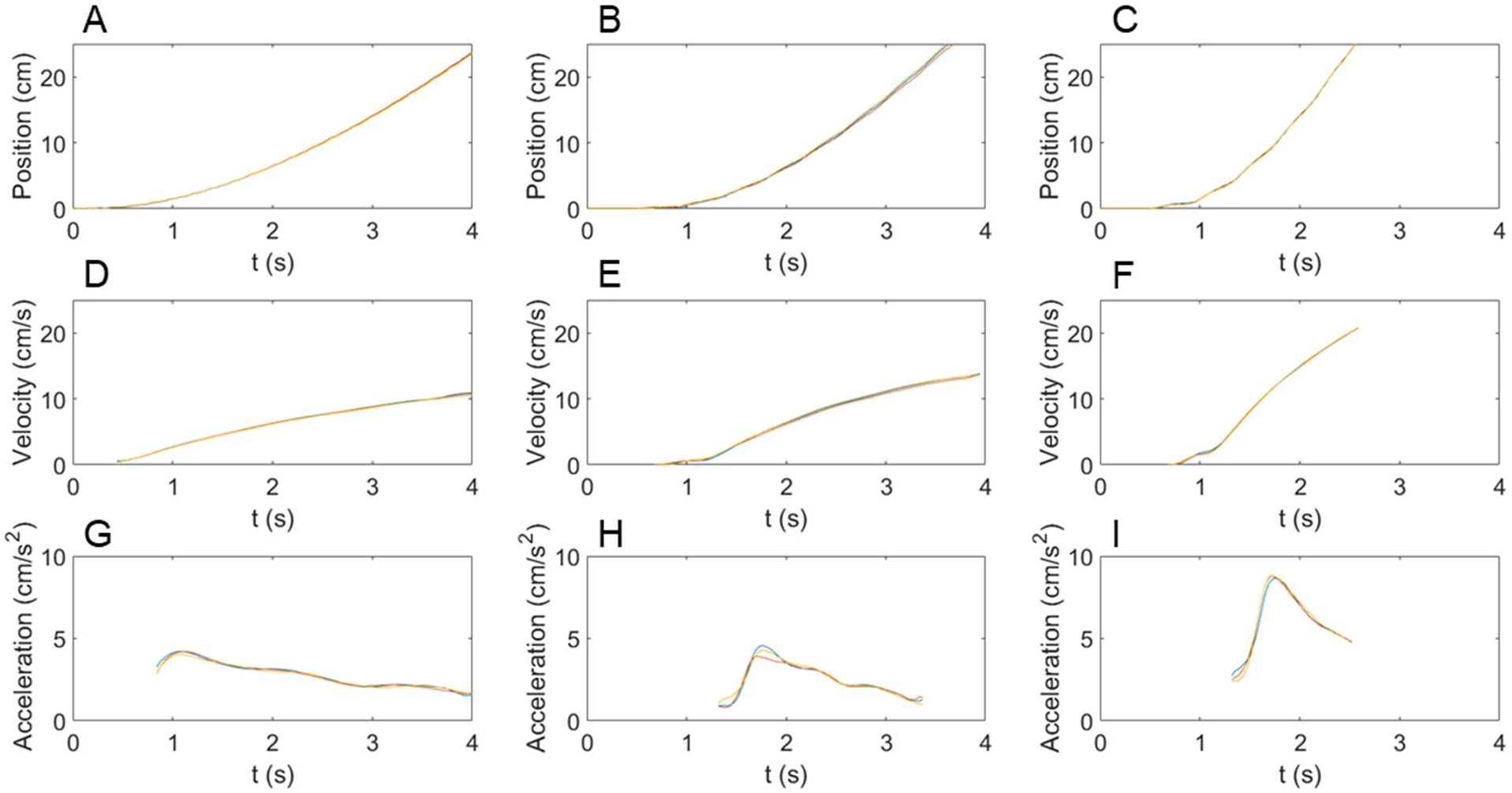
Representative examples of time-varying position, velocity, and acceleration data for the robotic model. The three lines within each plot represent independent trials under identical test conditions. (A, D, G) pure metachronal kinematics with *θ* = 75°, *ϕ* = 0%. (B, E, H) hybrid metachronal kinematics with *θ* = 75°, *ϕ* = 15%. (C, F, I) hybrid metachronal kinematics. with *θ* = 115° and *ϕ* = 15%. Position (A-C) and velocity (D-F) increased throughout the recording, while acceleration (G-I) reached a peak after a few cycles and then decreased. The peak value of the acceleration and the final value of the velocity were recorded and are reported in **Figure 5**.

## References

Alben S, Spears K, Garth S, Murphy D, Yen J (2010) Coordination of multiple appendages in drag-based swimming. J R Soc Interface 7(52), 1545–1557

Alexander DE (1988) Kinematics of swimming in two species of *Idotea* (Isopods: Valvifera). J Exp Biol 138(1), 37–49

Blake RW (1979) The mechanics of labriform locomotion I. labriform locomotion in the Angelfish *(Pterophyllum eimekei):* an analysis of the power stroke. J Exp Biol 82, 255–271

Campos EO, Vilhena D, Caldwell RL (2012) Pleopod rowing is used to achieve high forward swimming speeds during the escape response of *Odontodactylus havanensis* (Stomatopoda). J Crustac Biol 32(2), 171–179

Catton KB, Webster DR, Kawaguchi S, Yen J (2011) The hydrodynamic disturbances of two species of krill: implications for aggregation structure. J Exp Biol 214(11), 1845–1856

Colin SP, Costello JH, Sutherland KR, Gemmell BJ, Dabiri JO, Du Clos KT (2020) The role of suction thrust in the metachronal paddles of swimming invertebrates. Sci Rep 10(1), 1–8

Ford MP, Lai HK, Samaee M, Santhanakrishnan A (2019) Hydrodynamics of metachronal paddling: effects of varying Reynolds number and phase lag. R Soc Open Sci 6, 191387

Ford MP, Santhanakrishnan A (2021a) On the role of phase lag in multi-appendage metachronal swimming of euphausiids. Bioinspir Biomim, in press, doi: 10.1088/1748-3190/abc930

Ford MP, Santhanakrishnan A (2021b) Closer appendage spacing augments metachronal swimming speed by promoting tip vortex interactions. (Under Review)

Funfak A, Fisch C, Abdel Motaal HT, Diener J, Combettes L, Baroud CN, Dupuis-Williams P (2015) Paramecium swimming and ciliary beating patterns: a study on four RNA interface mutations. Integr Biol 7, 90–100

Granzier-Nakajima S, Guy RD, Zhang-Molina C (2020) A Numerical Study of Metachronal Propulsion at Low to Intermediate Reynolds Numbers. Fluids 5(2), 86

Hedrick TL (2008) Software techniques for two- and three-dimensional kinematic measurements of biological and biomimetic systems. Bioinspir Biomim 3(3), 034001

Heimbichner Goebel WL, Colin SP, Costello JH, Gemmell BJ, Sutherland KR (2020) Scaling of ctenes and consequences for swimming performance in the ctenophore *Pleurobrachia bachei*. Invertebr Biol 139(3)

Kiørboe T, Andersen A, Langlois VJ, Jakobsen HH (2010) Unsteady motion: escape jumps in planktonic copepods, their kinematics and energetics. J R Soc Interface 7(52), 1591–1602

Lim JL, DeMont ME (2009) Kinematics, hydrodynamics and force production of pleopods suggest jet-assisted walking in the American lobster *(Homarus americanus)*. J Exp Biol 212(17), 2731–2745

Murphy DW, Webster DR, Kawaguchi S, King R, Yen J (2011) Metachronal swimming in Antarctic krill: gait kinematics and system design. Mar Biol 158(11), 2541–2554

Murphy DW, Webster DR, Yen J (2012). A high-speed tomographic PIV system for measuring zooplanktonic flow. Limnol. Oceanogr. Methods 10(12), 1096–1112

Murphy DW, Webster DR, Yen J (2013) The hydrodynamics of hovering in Antarctic krill. Limnol Oceanogr Fluids Environ 3(1), 240–255

Murphy DW, Olsen D, Kanagawa M, King R, Kawaguchi S, Osborn J, Webster DR, Yen J (2019) The three-dimensional spatial organization of Antarctic krill schools. Scientific Reports 9, 381

Schabes M, Hamner W (1992) Mysid locomotion and feeding: kinematics and water-flow patterns of *Antarctomysis* sp., *Acanthomysis sculpta*, and *Neomysis rayii*. J Crustac Biol 12(1), 1–10

Schneider CA, Rasband WS, Eliceiri KW (2012) NIH Image to ImageJ: 25 years of image analysis. Nature Methods 9. 671–675

Takagi D (2015) Swimming with stiff legs at low Reynolds number. Phys Rev E 92(2), 023020

van Duren LA, Videler JJ (2003) Escape from viscosity: The kinematics and hydrodynamics of copepod foraging and escape swimming. J Exp Biol 206(2), 269–279

Walker JA (2002). Functional morphology and virtual models: physical constraints on the design of oscillating wings, fins, legs, and feet at intermediate Reynolds Numbers. Integ. And Comp. Biol. 42(2), 232–242

Wu JC (1980) Theory for aerodynamic force and moment in viscous flows. AIAA Journal 19(4). 432–441

Yen J, Brown J, Webster DR (2003) Analysis of the flow field of the krill, *Euphausia pacifica*. Mar Fresh Behav Physiol 36(4), 307–319

Zhang C, Guy RD, Mulloney B, Zhang Q, Lewis TJ (2014) Neural mechanism of optimal limb coordination in crustacean swimming. Proc Natl Acad Sci USA 111(38), 13840–13845

